# A20’s Linear Ubiquitin Binding Motif Restrains Pathogenic Activation of TH17/22 cells and IL-22 Driven Enteritis

**DOI:** 10.1101/2024.12.31.630926

**Authors:** Christopher John Bowman, Dorothea Stibor, Xiaofei Sun, Hiromichi Shimizu, Emily Yamashita, Nika Lenci, Rommel Advincula, Min Cheol Kim, Jessie A. Turnbaugh, Yang Sun, Bahram Razani, Peter J. Turnbaugh, Chun Jimmie Ye, Barbara A. Malynn, Averil Ma

**Author notes:** co-first author (Christopher John Bowman, Dorothea Stibor, Xiaofei Sun). Corresponding Address: Averil Ma, MD, 513 Parnassus Avenue, Medical Sciences Building, S-1032C San Francisco, California 94143, EM.

## Abstract

A20, encoded by the *TNFAIP3* gene, is a protein linked to Crohn’s disease and celiac disease in humans. We now find that mice expressing point mutations in A20’s M1 ubiquitin binding motif (ZF7) spontaneously develop proximate enteritis that requires both luminal microbes and T cells. Cellular and transcriptomic profiling reveal expansion of TH17/22 cells and aberrant expression of IL-17A and IL-22 in intestinal lamina propria of A20^ZF7^ mice. While deletion of IL-17A from A20^ZF7/ZF7^ mice exacerbates enteritis, deletion of IL-22 abrogates intestinal epithelial cell hyperproliferation, barrier dysfunction, and alarmin expression. A20^ZF7/ZF7^ TH17/22 cells autonomously express more RORγt and IL-22 after differentiation *in vitro*. ATAC sequencing identified an enhancer region upstream of the *Il22* gene in A20^ZF7/ZF7^ T cells, and this enhancer demonstrated increased activating histone acetylation coupled with exaggerated *Il22* transcription. Finally, CRISPR/Cas9-mediated ablation of A20^ZF7^ in human T cells increases RORγt expression and *IL22* transcription. These studies link A20’s M1 ubiquitin binding function with RORγt expression, epigenetic activation of TH17/22 cells, and IL-22 driven enteritis.

## Introduction

Intestinal immune homeostasis involves complex interactions between immune and non-immune cells. Many of these interactions are mediated by cytokines, and disruption of this cytokine network can lead to intestinal disease. Type 3 cytokines such as IL-17A, IL-17F, and IL-22 support anti-fungal and anti-bacterial responses (1). These mediators can also support epithelial homeostasis and regeneration (2). Dysregulated expression of type 3 cytokines is a feature of intestinal inflammation, and expansion of TH17 cells has been consistently observed in experimental models and human inflammatory bowel disease (IBD) patients (3, 4). However, IL-17A appears to play more complex roles in the intestinal milieu than in other tissues (5, 6). Some of this complexity may be related to the ability of TH17 cells to produce other cytokines under distinct physiological conditions. Hence, understanding the physiological regulation of intestinal cytokines and cytokine responses is crucial for dissecting the adaptive versus pathologic functions of these cytokine mediators.

A20/Tnfaip3 is genetically linked to IBD and celiac disease via GWAS (7–9). In addition, rare patients harboring haploinsufficient mutations of the A20 gene develop early onset IBD as well as Behcet’s disease with intestinal ulcerations (10–12). Hence, deficiencies of A20 expression and/or function likely compromise human intestinal homeostasis. Mechanistic studies have revealed that the A20 protein regulates several signaling cascades, including TNFR-, TLR-, TCR-, NOD2-, and CD40R-triggered signals (13–19). A20 performs these functions by regulating the ubiquitination of critical signaling molecules such as RIP1, RIP2, pro-IL-1β, and RIP3 as well as ubiquitinated signaling complexes such as the IKKγ complex. Yet, the mechanisms by which A20 preserves intestinal immunity are incompletely understood. This is partly because the A20 protein harbors distinct biochemical domains that mediate de-ubiquitinating (DUB), E3 ubiquitin (Ub) ligase, and non-catalytic linear (M1)-Ub chain binding activities (13, 14, 20–25). In this study, we unveil a unique role for A20’s M1-Ub binding motif in restraining epigenetic regulation of IL-22 expression in TH17/22 cells, intestinal epithelial cell homeostasis, and microbe-dependent enteritis.

## Results

### Linear (M1)-ubiquitin binding motif of A20 prevents T cell dependent enteritis

To understand the biochemical functions of A20 that regulate intestinal homeostasis, we analyzed intestines from a series of A20 knock-in mice that abrogate A20’s DUB (A20^OTU^), E3 Ub ligase (A20^ZF4^), or M1-Ub binding (A20^ZF7^) activities (20, 21). Macroscopically, small intestines from 12-week-old A20^ZF7/ZF7^ mice, but not other genotypes, demonstrated visible thickening of the intestinal wall. Histology revealed that the proximal small intestinal mucosa of A20^ZF7/ZF7^ mice harbored an increased accumulation of immune cells when compared with congenic A20^ZF4/ZF4^, A20^OTU/OTU^, or wildtype (WT) mice (**Figs. 1A,B**). As lymphocytes normally reside in the lamina propria, the intestines of A20^ZF7/ZF7^ mice contained plasma cells but, also, an expanded population of small lymphocytes. Granulocytes, including neutrophils, were absent. Overlying epithelium showed reactive changes, including goblet cell loss. Altogether, the expansion of the small intestine lamina propria (SILP) by lymphocytes is reminiscent of chronic enteritis, however other features of chronicity (eg, epithelial metaplasia, villus blunting) were absent. This phenotype was evident in both male and female A20^ZF7/ZF7^ mice and was 100% penetrant by 12 weeks of age. This phenotype was less pronounced in more distal portions of the small intestine, and large intestines from 12-week-old A20^ZF7/ZF7^ mice were not inflamed (**Supp. Fig. 1**). Hence, A20’s M1-Ub binding activity via its ZF7 domain preserves small intestinal immune homeostasis independently of A20’s DUB and E3 Ub ligase activities.

**Figure 1.**
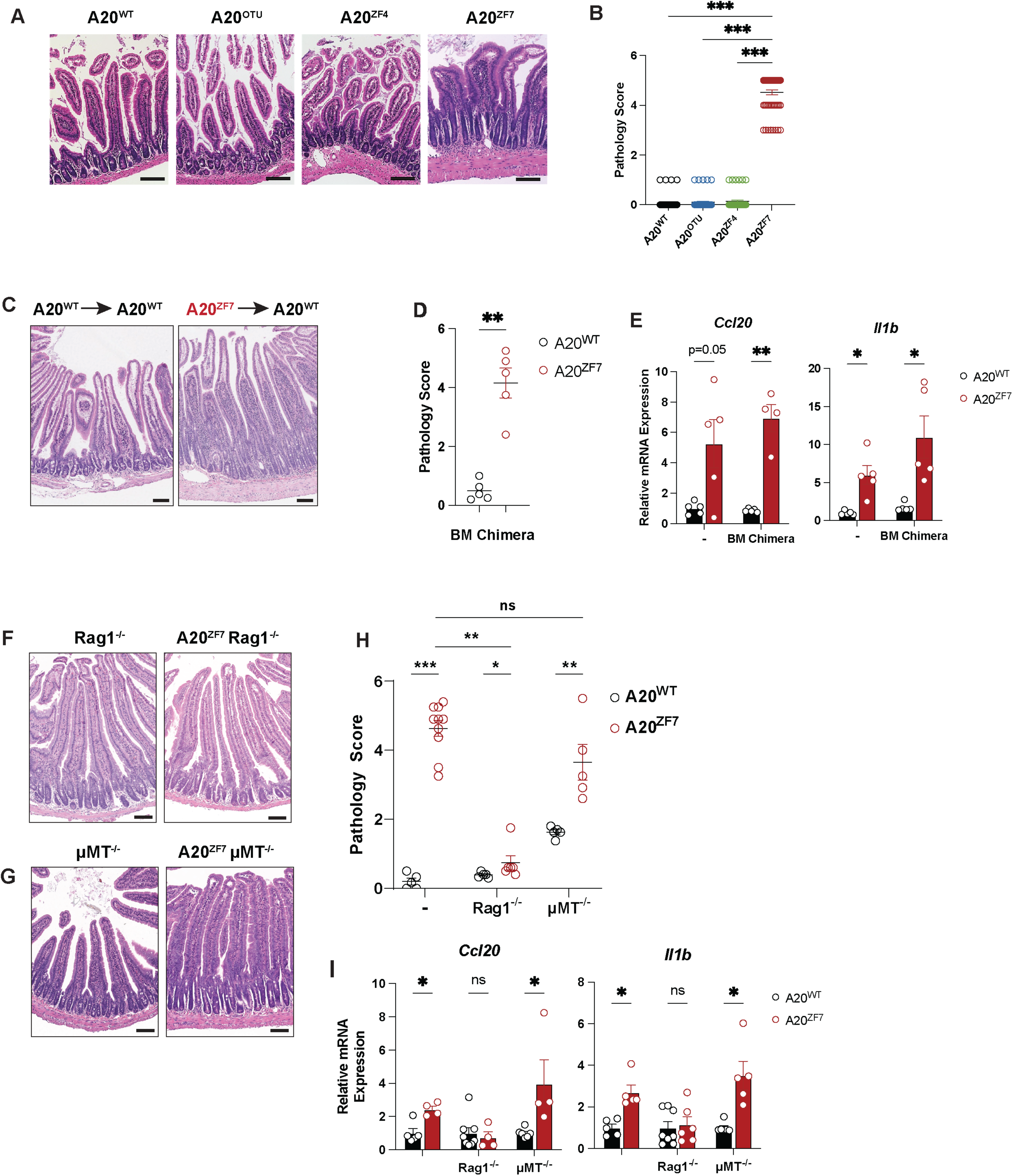
A20^ZF7^ restrains small intestinal enteritis that is T cell-dependent. (A, B) Representative H&E histology (A) and histopathological scores (B) of inflammation of proximate small intestines from indicated genotypes of mice. (Bar = 200 microns) (C, D) Representative H&E histology (C) and histopathological scores (D) of inflammation of small intestine from radiation bone marrow chimeric mice reconstituted with indicated bone marrow genotypes. (Bar = 100 microns.) (E) qPCR analyses of indicated pro-inflammatory genes in proximate small intestines from indicated bone marrow chimera. (F) Representative H&E histology of small intestine from indicated genotypes of RAG-1^-/-^ mice. Bar, 100 microns. Data representative from at least 4 independent pairs of mice. (G) Representative H&E histology of small intestine from indicated genotypes of μMT-null mice. Bar, 100 microns. Data representative from at least 3 independent pairs of mice. (H) Pathology scores of small intestinal inflammation. (I) qPCR analyses of indicated pro-inflammatory genes in proximate small intestines from indicated genotypes of mice. In figures above, each circle represents one mouse. Data shown as mean ± SEM. Statistics calculated using unpaired two-tailed Mann-Whitney U test (B, D, H) or unpaired two-tailed Student t-test with Welch correction (E, I). *p<0.05, ** p<0.01, ***p<0.001. ns, not significant.

A20 is expressed and induced in many cell types, including both hematopoietic and non-hematopoietic cells (26). To determine the degree to which A20^ZF7/ZF7^ mice (hereafter designated A20^ZF7^ mice) hematopoietic cells are sufficient to drive this pathology, we generated radiation chimera using either A20^ZF7^ or WT bone marrow cells to reconstitute irradiated WT mice. Chimera bearing A20^ZF7^ hematopoietic cells spontaneously developed enteritis by 12 weeks after reconstitution, a phenotype that was not seen in chimera containing WT hematopoietic cells (**Figs. 1C,D**). Furthermore, the A20^ZF7^ hematopoietic cells drive exaggerated expression of pro-inflammatory myeloid cytokines such as *Il1b* and chemokines such as *Ccl20* (**Fig. 1E**). To assess the relative contributions of T and B cells to enteritis in A20^ZF7^, we interbred these mice with RAG-1^-/-^ and μMT^-/-^ mice. Twelve-week-old A20^ZF7^ RAG-1^-/-^ mice exhibited negligible intestinal inflammation and expressed similar levels of *Il1b* and *Ccl20* as RAG-1^-/-^ mice (**Fig. 1F,H, I**), suggesting that adaptive lymphocytes are required for enteritis in A20^ZF7^ mice. By contrast, intestines from A20^ZF7^ μMT^-/-^ mice accumulated similar numbers of lamina propria immune cells (**Fig. 1G,H**) and expressed similarly elevated levels of inflammatory markers as A20^ZF7^ mice (**Fig. 1I**). Hence, RAG-1-dependent T lymphocytes, but not B lymphocytes, are required for small intestinal inflammation in A20^ZF7^ mice.

### Type 3 cytokines and TH17 cells are increased in intestinal lamina propria of A20^ZF7^ mice

To better understand why small intestinal inflammation develops in A20^ZF7^ mice, we analyzed the transcriptomes of intact small intestines from 12-week-old WT and A20^ZF7^ mice by bulk RNA sequencing. These studies revealed that A20^ZF7^ mice upregulated genes involved in NFκB and inflammasome signaling pathways (**Fig 2A**). In addition, A20^ZF7^ intestines expressed elevated levels of genes associated with CD4 T cell activation, T cell proliferation, and IL-1β production (**Fig. 2A**). IL-17 response genes were among the most enriched groups of genes, which – together with STAT and IL-6 regulation – suggests a prominent type 3 cytokine tone in A20^ZF7^ intestines. Indeed, quantitative mRNA analyses of intact intestines confirmed the exaggerated expression of type 3 cytokines, *Il17a* and *Il22* (**Fig 2B**). In addition, IL-22-dependent transcripts, such as *Reg3b*, *Reg3g*, *Saa1*, and *Saa3*, were significantly upregulated in A20^ZF7^ intestines (**Fig 2C, 5E**). As these data suggest that TH17 cells and/or group 3 innate lymphoid cells (ILC3s) might be expanded or hyperfunctional in A20^ZF7^ intestines, we profiled cellular infiltrates from small intestinal tissues. Immunohistochemistry of intact small intestines indicated CD4 T cells were expanded in the lamina propria of A20^ZF7^ mice (**Fig. 2D**), and flow cytometry of dissociated small intestinal lamina propria (SILP) cells confirmed a relative expansion of CD4 T cells (**Fig. 2E**). The increased number of CD4 T cells in A20^ZF7^ SILP largely comprised an expansion of TH17 cells when compared to WT littermates (**Fig. 2E**). In addition, among expanded A20^ZF7^ SILP CD4 T cells, IL-17A- and IL-17F-expressing cells were disproportionally increased while IFNγ expressing cells were present in similar proportions to WT mice.

**Figure 2.**
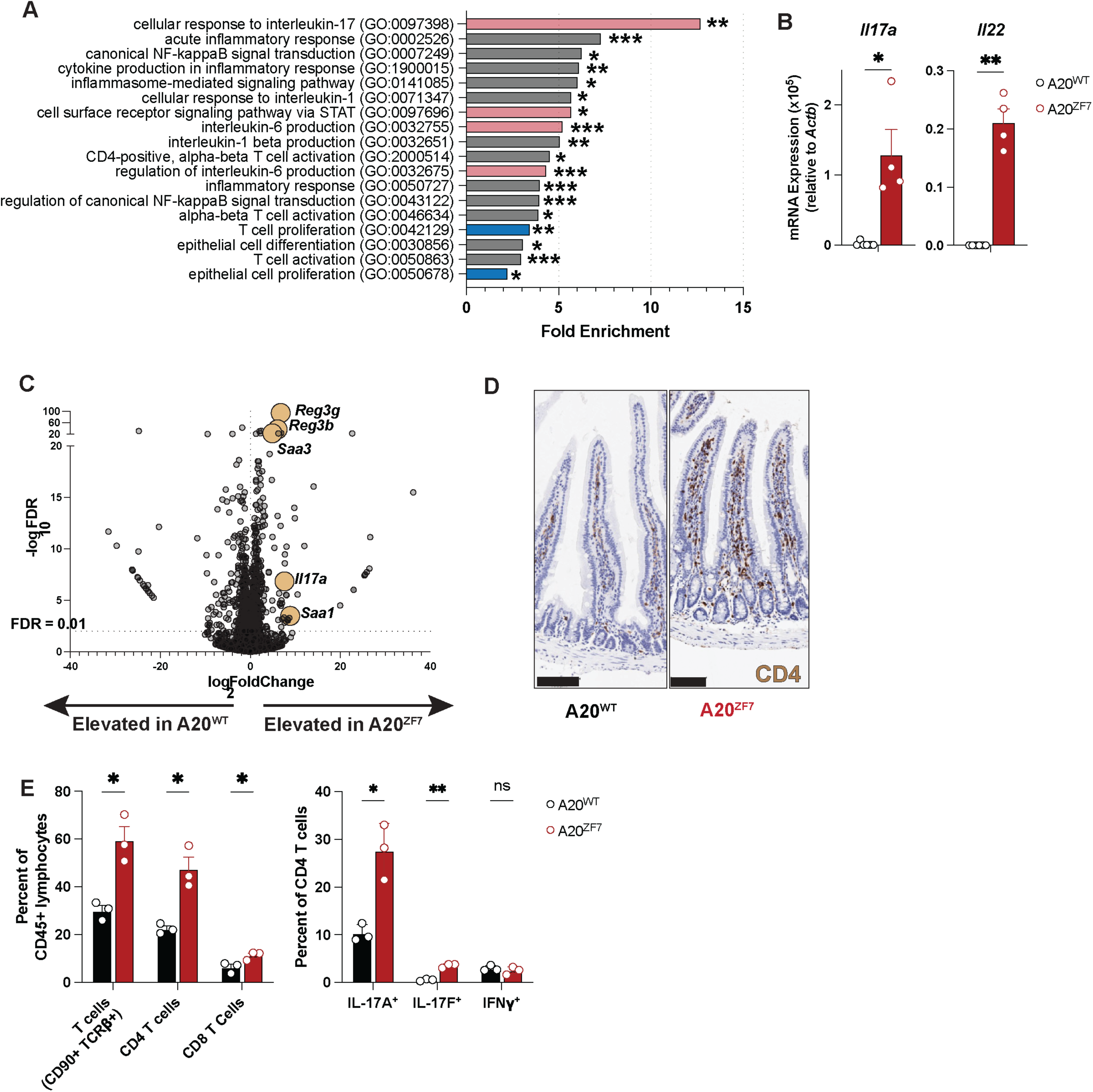
Small intestines of A20^ZF7^ mice harbor expanded TH17 cells. (A) Gene ontology enrichment of genes that are significantly upregulated in A20^ZF7^ versus WT intestines by bulk RNAseq. Red bars highlight categories related to TH17 differentiation. Blue bars highlight categories related to cellular proliferation. (B) qPCR analyses of *Il17a* and *Il22* expression from intact small intestine (relative to *Actb*). (C) Volcano plot of all annotated UCSC Refseq genes from bulk RNAseq analyses of A20^ZF7^ versus WT small intestines. Horizontal dashed line indicates adjusted p-value (FDR) of 0.01. (D) Representative immunohistochemical analyses of CD4 expression in WT and A20^ZF7^ small intestines. Data are representative of 3 mice from each genotype. Bar, 100 microns. (E) Flow cytometry of small intestinal lamina propria cells from WT and A20^ZF7^ mice. Data shown as mean ± SEM. Statistics calculated using unpaired two-tailed Student t-test with Welch correction. *p<0.05, ** p<0.01, ***p<0.001. ns, not significant.

To better define the SILP lymphocytes in A20^ZF7^ mice, we enriched SILP T cells from 12-week-old WT and A20^ZF7^ mice and profiled these cells by single cell RNA sequencing (scRNAseq). To avoid clustering cells based on cell cycle phases, genes related to cell cycling were removed prior to clustering analyses. UMAP analyses of these cell cycle-regressed cells identified CD8, TH17, and regulatory T cells, as well as subsets of innate lymphoid cells (ILCs) (**Figs 3A; Supp Fig 2A).** Genotype-specific UMAP analyses showed a relative expansion of CD4 and, to a lesser extent, CD8 T cells in A20^ZF7^ mice (**Figs 3B, C**). Expanded clusters of CD4 T cells in A20^ZF7^ intestines contained many cells expressing *Il17a* and *Il22*, delineating these cells as TH17 cells (**Fig. 3D; Supp Fig 2A)**. Another expanded cluster included regulatory T cells (**Fig 3C**). By contrast, group 3 ILCs (ILC3s) were relatively reduced in A20^ZF7^ mice (**Figs 3C).** In addition to being more abundant in A20^ZF7^ mice, A20^ZF7^ TH17 cells expressed higher levels of *Il17a* and *Il22* than WT TH17 cells (**Fig. 3E**). The marked expansion of TH17 cells, the increased expression of type 3 cytokines, and the absence of ILC3 expansion in A20^ZF7^ intestines relative to WT intestines suggest that TH17 cells account for the great majority of the increased type 3 cytokine production in A20^ZF7^ intestines.

**Figure 3.**
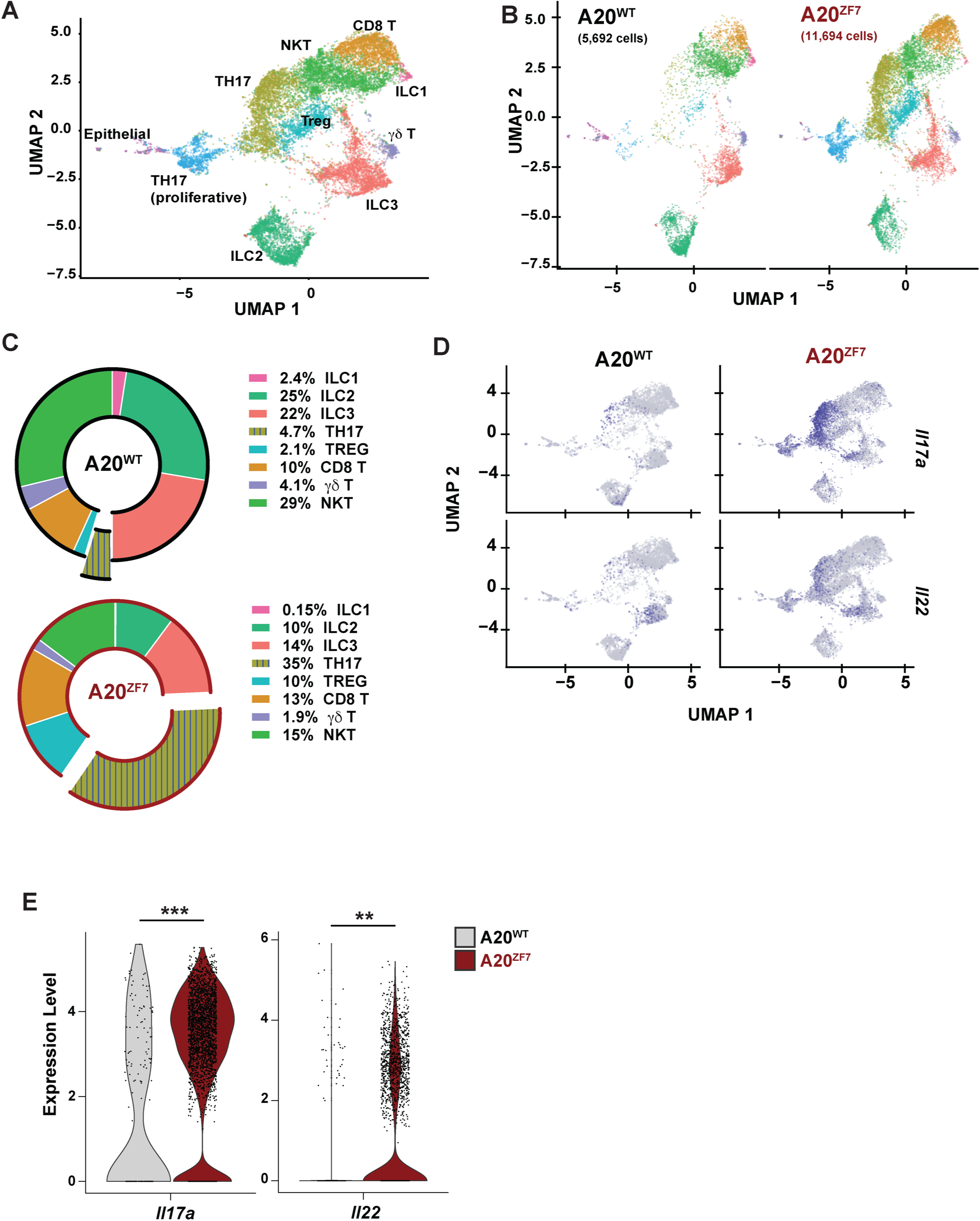
A20^ZF7^ mice show proliferation and expansion of TH17 and regulatory T cells within the small intestinal lamina propria (SILP) (A, B) UMAP clusters of scRNAseq analyses of SILP from WT and A20^ZF7^ mice. (C) Relative proportions of cell subsets in WT and A20^ZF7^ SILP. TH17 subset includes both proliferative (mustard yellow) and non-proliferative (sky blue) compartments. (D) Projection of *Il17a* and *Il22* expression onto UMAP clusters shown in panels A and B. (E) Violin plots of *Il17a* and *Il22* expression in TH17 cells from indicated genotypes of mice. Statistics calculated using unpaired two-tailed Wilcoxon rank sum test. *p<0.05, ** p<0.01, ***p<0.001. ns, not significant.

The TH17 cells segregated into two discrete clusters in UMAP space, and both clusters were heavily represented in A20^ZF7^ intestines (**Fig 3A,B; Supp Fig 2A**). Despite removing the contributions of cell cycle genes from affecting cell clustering, these two clusters exhibited differential cell cycle states: cells with high G0/G1 scores (indicating a predominantly non-proliferative state) and cells enriched for high G2/M or S scores (suggesting T cell proliferation) (**Supp Fig 2B**). In addition, this “TH17 proliferative” cluster expressed high levels of *Mki67*, the gene that encodes the proliferative marker Ki67 (**Supp Fig 2A**). The accumulation of proliferative TH17 cells in A20^ZF7^ SILP aligns with our bulk RNAseq analysis, which highlighted T cell proliferation as an enriched category in A20^ZF7^ intestines (**Fig 2A**). Although the proliferative cluster is transcriptionally distinct from non-proliferating T cells in either A20^ZF7^ or WT mice, it consists of cells that express detectable levels of *Il17a* and/or *Il22*, confirming they are TH17/22 cells (**Fig 3D; Supp Fig 2A**). Hence, increased numbers of proliferating TH17 cells helps explain why TH17 cells are more numerous in A20^ZF7^ intestines.

### IL-22, but not IL-17A, drives enteritis in A20^ZF7^ mice

As A20^ZF7^ small intestines contain dramatic expansions of SILP TH17 cells and increased tissue-wide expression of IL-17A and IL-22, we next sought to functionally define the potential roles of these cytokines in regulating intestinal disease in A20^ZF7^ mice. We interbred A20^ZF7^ mice with IL-17A^-/-^ mice and IL-22^-/-^ mice. Small intestines from A20^ZF7^ IL-17A^-/-^ mice exhibited more (rather than less) severe enteritis than IL-17A competent A20^ZF7^ mice (**Fig 4A, B**). Hence, IL-17A plays a protective rather than pro-inflammatory role in enteritis in A20^ZF7^ mice. A20^ZF7^ IL-17A^-/-^ intestines express more *Il22* than IL-17A^-/-^ but not A20^ZF7^ intestines (**Fig. 4C**). In contrast to A20^ZF7^ IL-17A^-/-^ mice, A20^ZF7^ IL-22^-/-^ mice exhibited less severe enteritis than A20^ZF7^ mice (**Fig. 4A, B**). Therefore, IL-22 promotes small intestinal inflammation in A20^ZF7^ mice. To better define the role of IL-22 in A20^ZF7^ intestines, we profiled the genome-wide transcriptomes of small intestines from A20^ZF7^ IL-22^-/-^ and control mice by bulk RNAseq. Principal component analysis (PCA) of bulk RNA-seq data revealed broad normalization toward wild-type transcriptomic states in A20^ZF7^ IL-22^-/-^ mice when compared with A20^ZF7^ mice (**Fig 4D**). Thus, IL-22 drives transcriptome-wide changes in the proximal small intestine to promote intestinal inflammation in A20^ZF7^ mice.

**Figure 4.**
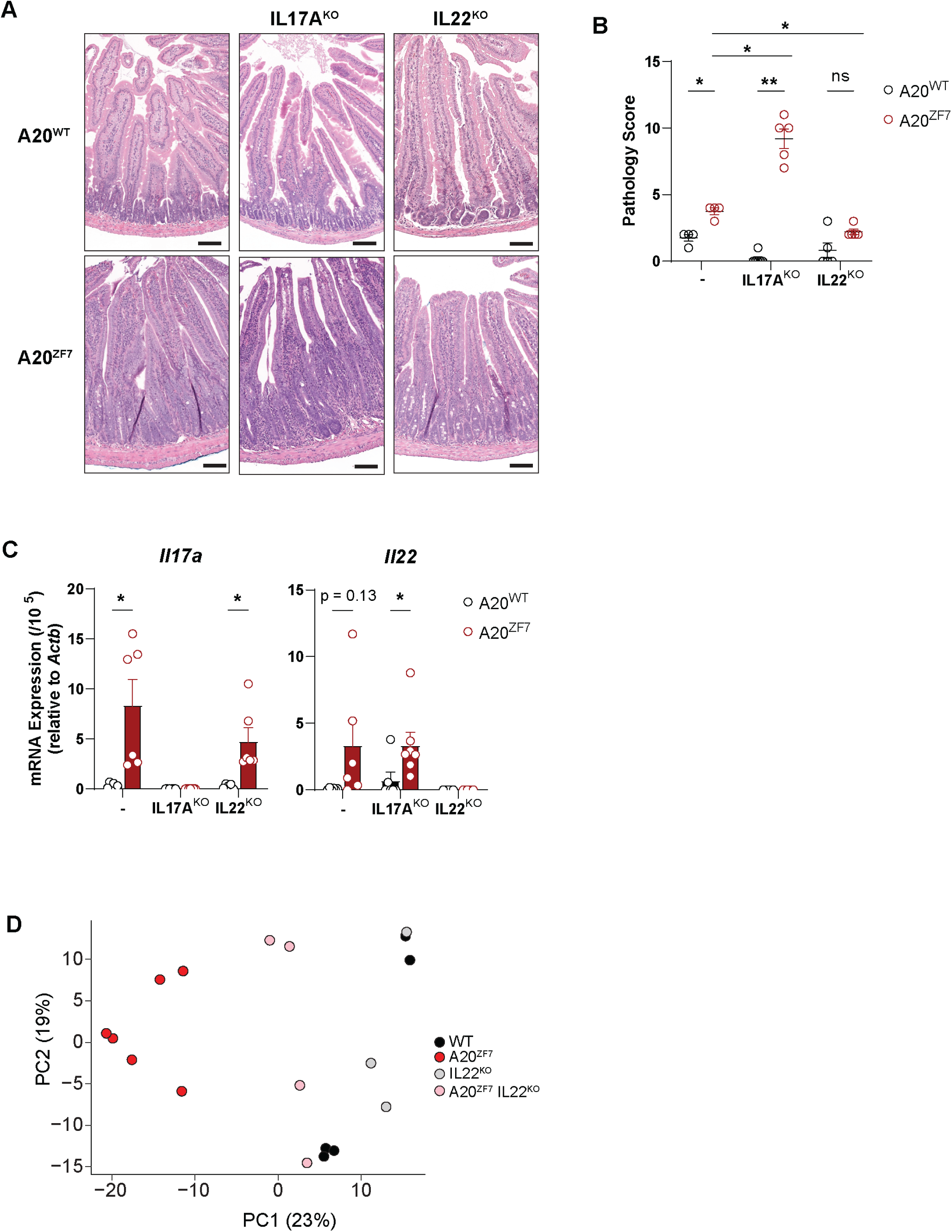
IL-17A protects against and IL-22 promotes small intestinal enteritis in A20^ZF7^ mice. (A, B) Representative H&E histology (A) and pathology scores (B) of small intestines from mice of the indicated genotypes. Bar= 100 um. (C) qPCR analyses of type 3 cytokines in small intestines from indicated genotypes of mice. (D) Principal component analyses of bulk RNAseq analyses of indicated genotypes of mice. Data shown as mean ± SEM. Statistics calculated using unpaired two-tail Mann-Whitney U test (B) or unpaired two-tailed Student t-test with Welch correction (C). *p<0.05, ** p<0.01, ***p<0.001. ns, not significant.

### A20^ZF7^ mice exhibit IL-22- and microbe-dependent epithelial barrier dysfunction

Il22ra1, the IL-22-specific receptor chain, is selectively expressed on intestinal epithelial cells (IEC) but not immune cells (27). Hence, pathophysiological effects of IL-22 in A20^ZF7^ mice could be mediated by perturbation of IEC functions. By histology, epithelial crypts were disproportionally expanded in small intestines of A20^ZF7^ mice when compared with WT mice (**Figs. 4A, 5A**). Bulk RNAseq highlighted epithelial cell proliferation as an enriched gene set in A20^ZF7^ intestines (**Fig 2A**), and immunohistochemistry for Ki67 confirmed an expansion of proliferating crypt IECs in A20^ZF7^ intestines (**Fig 5A,B**). Interestingly, chimera bearing A20^ZF7^ hematopoietic cells also expressed elevated levels of IEC-derived defensins *Reg3b* and *Reg3g* (**Fig 5C**). This result supports that radiation-sensitive A20^ZF7^ TH17 cells express exaggerated amounts of IL-22 that perturb WT IECs. Notably, the IEC proliferation and crypt elongation seen in A20^ZF7^ mice were normalized in A20^ZF7^ IL-22^KO^ mice (**Fig 5A,B**). To understand the IL-22-dependent perturbations of IECs in A20^ZF7^ mice, we isolated small intestinal IECs from A20^ZF7^ IL-22^KO^ and control mice. Transcriptomic analyses of these cells revealed that A20^ZF7^ epithelia expressed more C-type lectins (*Reg3b*, *Reg3g*), alarmins (*Saa1*), and chemokines (*Cxcl1*) than WT IECs (**Fig. 5D,E**). These defects were markedly reduced in A20^ZF7^ IL-22^KO^ IECs, indicating that IL-22 drives exaggerated expression of these pro-inflammatory mediators in A20^ZF7^ IECs.

**Figure 5.**
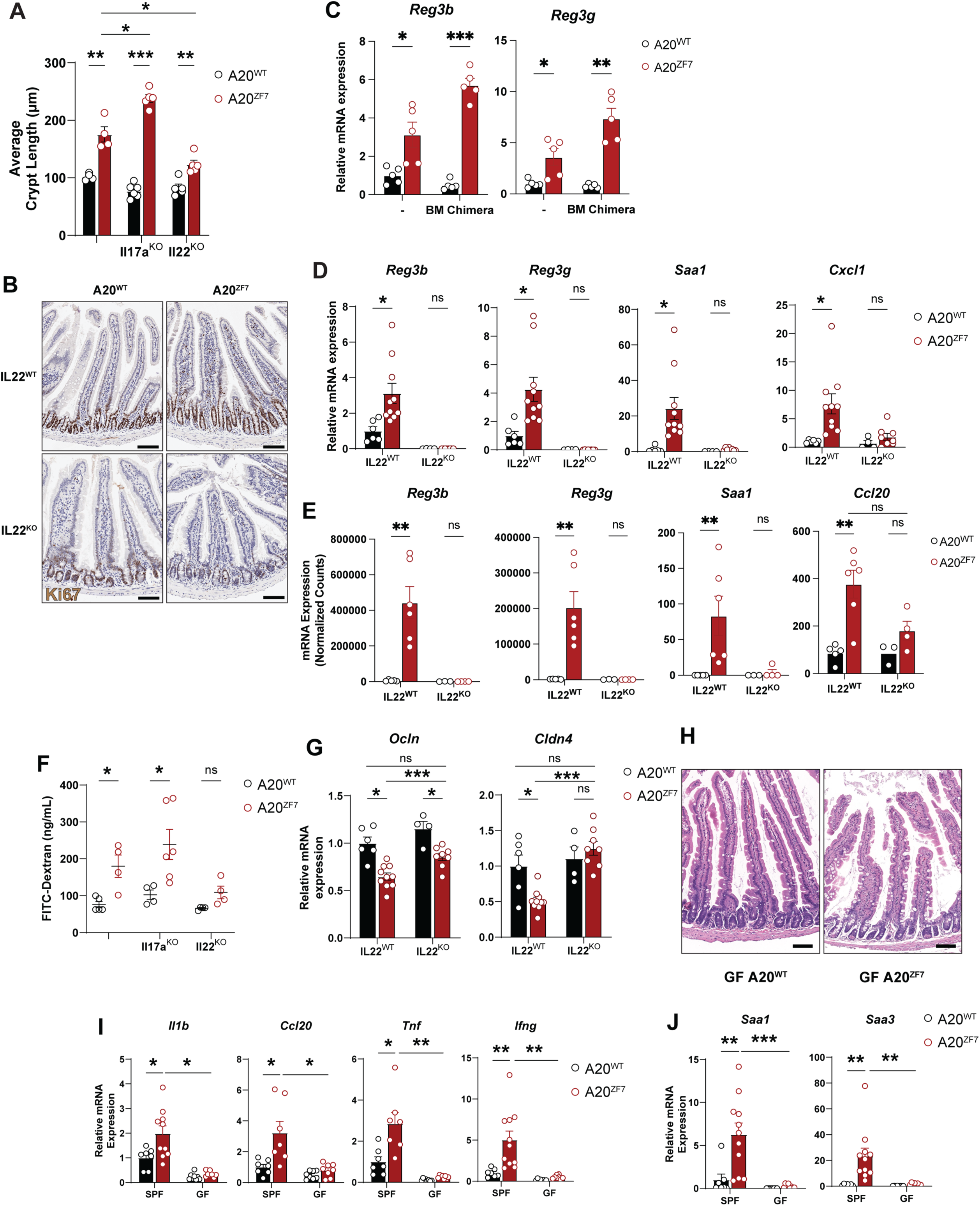
IL-22 disrupts epithelial barrier integrity and drives microbe-dependent enteritis in A20^ZF7^ mice. (A) Average crypt length measured in histology sections of proximal small intestines from indicated genotypes of mice. (B) Representative immunohistochemistry of Ki67 expression in small intestines of indicated genotypes of mice. Data are representative of 2-3 mice from each genotype. Bar= 100 um. (C) qPCR analyses of IL-22 dependent defensins from intact small intestines of indicated radiation chimera. (D) qPCR analyses of IL-22 dependent genes from isolated small intestinal IECs. (E) Normalized mRNA counts of IL-22 dependent genes from bulk RNA-seq of intact small intestines. (F) Fluorescence measurements of FITC-dextran in sera from indicated genotypes of mice four hours after FITC-dextran gavage. (G) qPCR analyses of indicated tight-junction genes from isolated small intestinal IECs from indicated genotypes of mice. (H) Representative H&E histology of proximate small intestines from germ-free mice of indicated genotypes. Bar= 200 um. (I,J) qPCR analyses of indicated genes from small intestines from indicated genotypes of specific pathogen-free (SPF) and germ-free (GF) mice. Data above shown as mean ± SEM. Each circle represents an independent mouse. Statistics calculated using unpaired two-tailed Student t-test with Welch correction. *p<0.05, ** p<0.01, ***p<0.001. ns, not significant.

As pro-inflammatory cytokines can perturb epithelial barrier functions (28–30), we tested intestinal epithelial permeability in A20^ZF7^ IL-22^KO^ mice and control mice by gavaging these mice with FITC-dextran. Increased absorption of FITC-dextran in sera of A20^ZF7^ mice supports aberrant barrier integrity in these mice. This compromised barrier integrity is similarly exacerbated in A20^ZF7^ IL-17A^-/-^ mice but normalized in A20^ZF7^ IL-22^-/-^ mice (**Fig. 5F**). To understand why A20^ZF7^ IECs fail to maintain barrier integrity, we assayed expression of occludin (Ocln) and claudin-4 (Cldn4), two tight junction proteins that support epithelial barrier integrity (31, 32). *Ocln* and *Cldn4* expression were depressed in A20^ZF7^ compared to WT IECs and were normalized in A20^ZF7^ IL-22^KO^ IECs (**Fig. 5G**). Diminished barrier integrity may allow translocation of microbes and pathogenic products, inciting an inflammatory response (33–35). To determine whether intestinal florae are required for enteritis in A20^ZF7^ mice, these mice were derived into germ-free environments. Germ-free A20^ZF7^ mice exhibited neither intestinal inflammation (**Fig 5H**) nor elevated expression of proinflammatory cytokines (ie*, Il1b, Ccl20, Tnf, Ifng*) relative to controls (**Fig 5I**), a stark contrast to the A20^ZF7^ mice raised in specific pathogen-free environments. Moreover, despite the presence of A20^ZF7^ in the small intestines of germ-free mice, the absence of commensal microbiota did not lead to alarmin (*Saa1, Saa3*) production (**Fig 5J**), suggesting that microbiota are necessary for IL-22-dependent programs within the small intestine. Therefore, IL-22 and microbes drive enteritis in A20^ZF7^ mice that is associated with disruption of intestinal epithelial barrier integrity and homeostasis.

### A20^ZF7^ restrains IL-22 expression in murine and human CD4^+^ T cells

Our data above show CD4 T cell autonomous A20^ZF7^ functions and IL-22 are integral to enteritis in A20^ZF7^ mice. Accordingly, we investigated how A20^ZF7^ regulates T cell production of IL-22. We enriched naïve CD4^+^ T cells from 8-week-old A20^ZF7^ and WT mice and differentiated these cells into TH17 cells with recombinant IL-6 and TGFβ. The efficiency of *in vitro* TH17 differentiation (as defined by the expression of RORγt, the key transcription factor that coordinates TH17 differentiation (36) was consistently >99% (**Fig 6A**), regardless of A20 mutation status. Notably, A20^ZF7^ TH17 cells expressed RORγt in higher amounts than WT TH17 cells, suggesting that A20^ZF7^ restrains RORγt expression in a cell autonomous fashion (**Fig 6A**). In the absence of PMA/ionomycin stimulation, A20^ZF7^ TH17 cells also expressed more *Il22* mRNA than WT cells (**Fig. 6B**). Since *Il22* expression is dependent on the aryl hydrocarbon receptor (Ahr) (37), these cells were treated with the Ahr agonist FICZ (37, 38). FICZ further exaggerated enhanced *Il22* expression in A20^ZF7^ TH17 cells relative to WT cells (**Fig 6B**). Similarly, ELISA of supernatants from these cells revealed that A20^ZF7^ T cells secreted more IL-22 protein than WT cells (**Fig. 6C**). Notably, these results were obtained from cells that were not stimulated with PMA or ionomycin, avoiding potential caveats associated with possible A20-dependent regulation of these stimuli. Aligned with these findings, flow cytometry studies of PMA/ionomycin-stimulated cells showed increased numbers of IL-22^+^ T cells in A20^ZF7^ cultures when compared to WT cultures (**Supp Fig. 3B**). Hence, mutation of A20^ZF7^ within T cells leads to greater RORγt expression and increased expression of IL-22.

**Figure 6.**
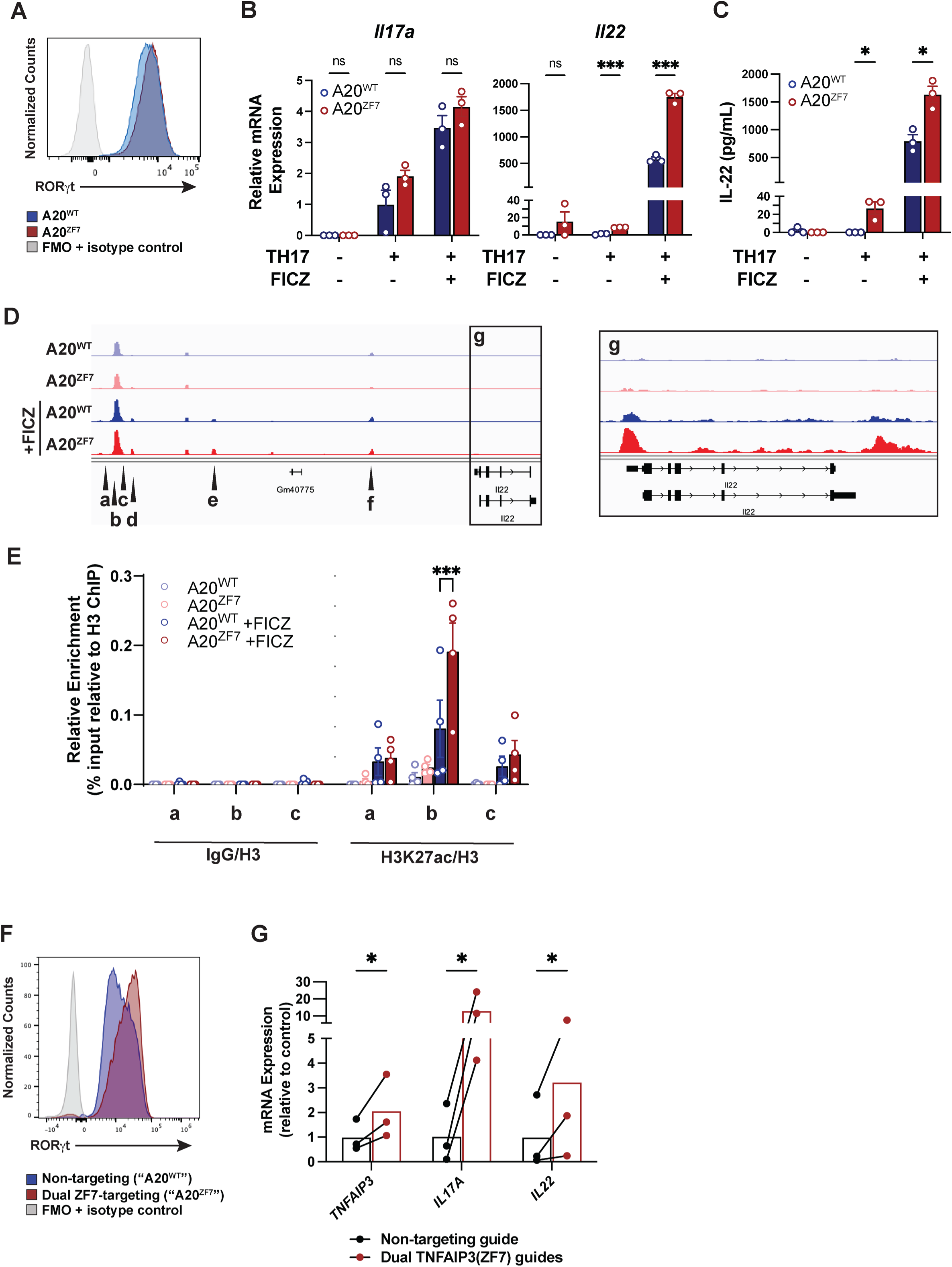
A20^ZF7^ TH17 cells show enhanced IL-22 production due to increased histone acetylation at an *Il22* enhancer. (A) Flow cytometric analysis of RORγt expression in *in vitro* differentiated TH17 cells from WT and A20^ZF7^ mice. (B) qPCR analyses of *Il17a* and *Il22* expression in cells generated as in (A) above. “TH17” indicates TH17 differentiation conditions. “FICZ” indicates treatment with the AHR agonist FICZ. Note increased IL-22 expression in absence of supplementary treatment with PMA/ionomycin. (C) ELISA of IL-22 secretion from cells generated as in (A). (D) ATAC-seq of genomic loci at/near the *Il22* locus in cells generated as in (A) above. Note increased DNA accessibility across *Il22* gene in enlarged plot (locus g). (E) Chromatin IP of acetylated H3K27 at indicated *Il22* loci (a,b,c in (D) above) in cells generated as in (A) above. Note increased H3K27 acetylation at locus b in A20^ZF7^ cells, coinciding with increased *Il22* transcription. (F) Flow cytometric analyses of RORγt expression in CRISPR/Cas9-edited primary human T cells differentiated *in vitro* using TH17 conditions. (G) qPCR analyses of expression of indicated genes in paired isogenic human TH17 cells that were engineered with CRISPR/Cas9 and either A20^ZF7^ targeted or non-targeting guide RNAs. Note increased expression of *TNFAIP3*, *IL17A*, and *IL22* in A20^ZF7^ ablated TH17 cells. Three pairs of isogenic samples from two healthy donors are shown. Data shown as mean ± SEM. Statistics calculated using unpaired two-tailed Student t-test with Welch correction (B, C, E) or paired ratio t-test (G). *p<0.05, ** p<0.01, ***p<0.001. ns, not significant.

As A20^ZF7^ TH17 cells express more *Il22* mRNA and protein than WT cells, we hypothesized that A20^ZF7^ restrains epigenetic regulation of *Il22* transcription. To identify potential genomic loci that may regulate *Il22* in cis, we surveyed chromatin accessibility via ATAC sequencing on TH17/22 cells differentiated from naïve WT and A20^ZF7^ CD4 T cells. ATAC-seq revealed increased accessibility across the *Il22* gene in A20^ZF7^ TH17 cells (**Fig. 6D, region g**), supporting open chromatin at *Il22*’s promoter and enhanced *Il22* transcription. In addition, multiple other sites upstream of *Il22* demonstrated DNA accessibility, predicting potential enhancer regions (**Fig 6D, regions a-f**). Notably, ATAC-seq highlighted a conserved region 32 kb upstream of *Il22*’s transcription start site (**Fig. 6D, region b**), a site recently characterized as an *Il22* enhancer (39). To define this conserved region as a bona-fide functional enhancer and to quantify the activation state of this enhancer, naïve WT and A20^ZF7^ CD4 T cells were differentiated into TH17/22 cells, and chromatin immunoprecipitation (ChIP) of acetylated lysine 27 of histone H3 (H3K27ac) was performed at these genomic loci. H3K27ac was increased in A20^ZF7^ TH17 cells at the enhancer for *Il22* (**Fig. 6E, region b**), supporting increased enhancer activation and enhancement of *Il22* transcription in A20^ZF7^ TH17 cells. By contrast, sequences located on the immediate shoulders of the enhancer **(Fig 6E, regions a and c**) as well as other DNA accessible sites (**Supp Fig 3C, regions d-f**) were less decorated with H3K27ac and similarly marked in WT and A20^ZF7^ TH17 cells. The enhancement of H3K27ac marks at the conserved enhancer with FICZ treatment suggests Ahr activation facilitates the acetylation of this locus. These results provide a molecular underpinning for Ahr and FICZ’s roles in promoting *Il22* expression (37, 38). Taken together, these studies suggest that A20^ZF7^ restrains hyperacetylation of a specific *Il22* enhancer and *Il22* transcription in CD4 TH17 cells.

The *A20/TNFAIP3* gene is well-conserved between murine and human genomes, and A20 polymorphisms and mutations are associated with human IBD and celiac disease (7–10, 40). To determine if A20’s ZF7 domain regulates TH17 cells functions and IL-22 expression in human T cells, we generated A20^ZF7^ mutant human CD4 T cells using CRISPR/Cas9. Since A20’s ZF7 domain comprises the most C-terminal residues of the A20 protein, targeting A20^ZF7^ should not affect the stability or structure of the A20 protein. Our recent studies of murine A20^ZF7^ mutant proteins suggest that these proteins are indeed expressed at supranormal levels in TNF-stimulated fibroblasts (21). We thus designed CRISPR guide RNAs (gRNAs) to delete the N-terminal half of A20^ZF7^. We isolated naïve CD4 T cells from healthy donor peripheral blood, activated the cells via TCR stimulation, and electroporated CRISPR/Cas9 with either these ZF7-targeting or control gRNAs to generate A20^ZF7^ mutant or isogenic control human T cells, respectively. Sanger sequencing demonstrated >85% of alleles were targeted by A20^ZF7^-specific gRNAs (**Supp Fig 4A,B**). These cells were then differentiated toward TH17/22 cells for 9 days, at which time more than 99% expressed RORγt in both A20^ZF7^-ablated and control cells (**Fig 6F**), confirming efficient TH17/22 differentiation. Expression of *A20/Tnfaip3* mRNA is markedly induced by NFκB activity, and A20 mediates negative feedback on NFκB signaling (26). We observed increased expression of *TNFAIP3* mRNA in A20^ZF7^ human T cells when compared to paired isogenic control T cells (**Fig. 6G**). This result suggests that A20^ZF7^ T cells exhibit increased NFκB signaling. The NFκB family members c-Rel and Rela/p65 bind RORγt promoters and drive RORγt transcription (41). Consistent with increased NFκB signaling and with our findings in murine TH17 cells, ablation of A20^ZF7^ in human TH17 cells led to higher expression of RORγt (**Fig. 6F**). Finally, A20^ZF7^-deleted human TH17/22 cells led to elevated expression of *IL17A* and *IL22* transcripts relative to paired, isogenic, A20-competent control cells (**Fig. 6G**). Hence, A20’s ZF7 motif restrains NFκB activity, RORγt expression, and *IL22* expression in human TH17 cells. This function thus represents a new molecular lever regulating pathogenic activation of both murine and human TH17/22 cells.

## Discussion

Our studies uncover a new spontaneous model of proximate enteritis that links A20’s M1-Ub binding motif to pathogenic TH17 cell activation, IL-22-dependent epithelial dysfunction, microbe-dependent enteritis, and epigenetic regulation of *Il22* transcription. Inflammation of the proximate small bowel in A20^ZF7^ mice aligns with two human diseases that can afflict this portion of the intestine: Crohn’s disease and celiac disease. In addition, rare patients with monogenic A20 haploinsufficiency can develop very early onset IBD or Behcet’s disease with intestinal ulceration. All these diseases are genetically linked to A20 deficiency. We show that non-enzymatic, Ub binding by A20^ZF7^ is more important than ZF4’s Ub binding or E3 Ub ligase activity for preserving small intestinal homeostasis. The selective importance of ZF7’s Ub binding motif may be related to its preferential binding to M1-linked Ub dimers (23), its increased binding affinity to Ub tetramers (22), and/or its ability to bind and regulate IKKγ signaling complexes (21). Haploinsufficient HA20 patients harbor various *TNFAIP3* mutations, and a common deficit of these mutations is they cause premature stop codons that result in the loss of A20’s C-terminal ZF7 domain (10, 40, 42). Therefore, our findings not only highlight the importance of the non-enzymatic ZF7 functions of the A20 protein but also broadly identify a unifying molecular dysfunction that perpetuates intestinal disease in human patients.

Our studies provide new insights into the homeostasis of intestinal tissue-resident TH17 cells. Selective expansion of IL-17A and IL-22 expressing cells in A20^ZF7^ mice mirrors the expansion of pro-inflammatory TH17/22 cells observed in the intestines of human IBD patients (43). Expansion of TH17 cells is likely coupled with activation of these cells and a transition from a homeostatic state (wherein cells express IL-17A) to a pro-inflammatory state characterized by the additional expression of IL-17F, IL-22, GM-CSF, MCSF, and granzymes (44, 45). This important transition can be stimulated by IL-23 (44, 46). The importance of these TH17 cell-derived cytokines other than IL-17A in the intestine has been implicated by the anti-inflammatory effect of IL-23 blockade coupled with pro-inflammatory consequences of IL-17A inhibition (47). Our studies highlight the ability of IL-22 to drive proximate enteritis. Increased expression of other TH17 cell cytokines in A20^ZF7/ZF7^ intestines, e.g., IL-17F, IFNγ, may also contribute to enteritis in these mice, as IL-22 deficiency partially rescues this phenotype. More broadly, increased cytokine expression by A20^ZF7^ TH17 intestinal T cells implicates A20^ZF7^ as a critical mediator of TH17 cell activation and/or quiescence. Mutation of this biochemical motif likely leads to a failure of TH17 cells to properly regulate homeostatic TCR or cytokine receptor signals and prevent pathologic activation of these cells.

Our studies unveil new mechanisms by which IL-22 triggers inflammation in the bowel, including elevated levels of IEC alarmins. Among cytokines that directly regulate IEC functions, IL-22 is distinct from IL-17A, IL-17F, and IFNγ in that IL-22 binds to IECs and not immune cells. In this regard, IL-22 occupies a unique niche in immune-epithelial crosstalk. While prior studies showed that IL-22 supports reparative functions in IECs (2, 48, 49), IL-22 can also mediate inflammation when Treg- or IL-10-deficiency causes aberrant macrophage activation and colitis (50). By contrast, A20^ZF7/ZF7^ mice develop enteritis in the presence of supranormal levels of Tregs and IL-10. Hence, our studies reveal that IL-22 can cause intestinal inflammation despite intact Treg functions. IL-22-dependent pathophysiology in A20^ZF7/ZF7^ mice also appears distinct from that seen in Treg- and IL-10-deficient mice in that A20^ZF7/ZF7^ mice develop proximal enteritis, while the latter predominantly develop colitis. We have found that IL-22 stimulates exaggerated IEC elaboration of alarmins, hyperproliferation of crypt IECs, and compromised epithelial permeability, leading to microbe-dependent enteritis. These epithelial dysfunctions broaden IL-22-dependent biology beyond its tissue reparative and defensin activities. They demonstrate that dysregulated IL-22 expression from pathogenically activated TH17/22 cells can profoundly disrupt IEC homeostasis. The context-specific variables that influence whether IL-22 performs pro-inflammatory versus tissue reparative functions may include the cellular source of IL-22 (i.e., TH17/22 versus ILC3 cells), the epithelial subtypes (e.g., small versus large intestine; crypt progenitor vs mature epithelial cells), co-expressed cytokines, and/or the abundance or duration of IL-22 signals.

We have found that enteritis in A20^ZF7/ZF7^ mice is microbe-dependent. Yet this enteritis selectively afflicts the proximal small intestine that is typically colonized with fewer numbers of bacteria than more distal regions. Segmented Filamentous Bacteria and *Citrobacter rodentium* perturb intestinal immune homeostasis in the ileum and colons of mice, respectively, and this selectivity is likely related to the niches in which these organisms colonize the lumen. Hence, it is possible that microbes that selectively colonize the proximal small intestine may drive enteritis in this region of A20^ZF7/ZF7^ mice. Small intestinal microbes have recently been highlighted to alter lipid absorption (51). Alternatively, IECs in the proximal small intestine of these mice may harbor preferential sensitivity to microbially triggered IL-22 signals. Other potential contributing factors include luminal products that are concentrated in the proximal bowel, e.g., bile acids secreted into the lumen via the bile duct. As we have found that germ-free conditions abrogate enteritis, these moieties could be secondary bile acids that are modified by small intestinal bacteria. Future studies of duodenal microbes and/or bile acid metabolites could unveil mechanisms by which homeostasis is preserved in the duodenum.

We have uncovered T cell-autonomous functions for A20^ZF7^ that restrain pathogenic expression of IL-22. Prior work from our lab and others showed that A20 restrains TCR-induced NFκB signaling in naïve T cells (15, 18, 52). Hence, intestinal A20^ZF7/ZF7^ TH17 cells may exhibit NFκB signaling in response to homeostatic TCR signals. Notably, the enhanced *Il22* transcription we observe in murine A20^ZF7/ZF7^ TH17 cells *in vitro* occurs in the absence of IL-1 or IL-23. Prior studies showed that c-Rel was required for driving RORγt promoter activity (41) and, combined with our current work, suggests that increased NFκB activity in A20^ZF7/ZF7^ TH17 cells stimulates increased RORγt expression. We surmise that enhanced RORγt in A20^ZF7^ T cells stimulates TH17 differentiation and, in the context of A20 defects, may promote pathogenic activation of *Il22* transcription via an *Il22* enhancer that harbors NFκB, RORγt, Runx1, and AP-1 sites (39). Intriguingly, this murine enhancer is also highly conserved upstream of the human *IL22* locus. Combined with our finding that ablation of A20^ZF7^ in human T cells causes increased *IL22* expression, inhibition of this enhancer by A20^ZF7^ prevents pro-inflammatory expression of IL-22 and may restrain a broader program of pathogenic activation of both murine and human TH17 cells. Given the importance of pathogenically activated TH17 cells to intestinal inflammation, this biochemical function provides an important new lever for preserving TH17 cell quiescence and preventing human disease.

## Materials and Methods

### Sex as a Biological Variable

Both male and female mice were used in this study, and similar results were obtained with both. Sex is not considered a biological variable in this study.

### Mice

Animal studies were conducted under an approved Institutional Animal Care and Use Committee (IACUC) protocol at University of California, San Francisco (UCSF). Mice were bred and housed at 22°C under a 12 hr light:dark cycle with ad libitum access to food and water. A20^C103/C103^ (OTU), A20^ZF4/ZF4^ (ZF4), and A20^ZF7/ZF7^ (ZF7) knock-in mice were generated in our laboratory and previously described (20, 21). B6.129S2 Ighm^tm1Cgn^/J (μMT; 002288), B6.129S7 RAG-1^tm1Mom^/J (Rag1; 002216), and C57BL/6 Il22^tm1.1(icre)Stck^/J (IL22; 027524) mice were purchased from Jackson Laboratories. IL-17A^-/-^ mice were kindly provided by Y. Iwakura via S. Gaffen.

### Tissue histology and scoring

Mice at the indicated ages were euthanized, and the proximal small intestine was collected in 4% paraformaldehyde. Samples were subsequently processed and stained by HistoWiz to produce hematoxylin and eosin (H&E), CD4-, or Ki67-stained slides. Pathology scores of intestinal inflammation were generated by assessing total tissue inflammation and epithelial changes in the colon. The total inflammation score (scale 0-3) was based on the overall severity or extent of inflammation in the intestine. An epithelial change score (scale 0-4) was assigned for each of the following categories: goblet cell loss, intraepithelial neutrophils and/or cryptitis, abscesses, or crypt loss. The scores were summed to generate a pathology score for each mouse.

### Radiation Bone Marrow (BM) Chimera

At 6 weeks of age, wild-type recipient mice were irradiated with 1200 rads total body radiation. The same day, recipients were injected intravenously with 5 × 10^6^ BM cells isolated from femurs and tibias of corresponding donors (WT or A20^ZF7^ mice). Animals were kept on antibiotics in the drinking water for one week after irradiation. BM recipients were sacrificed 12 weeks after BM transfer.

### FITC-Dextran Epithelial Barrier Assay

Mice (11-week-old) were fasted for five hours prior to oral gavage of FITC-dextran (average MW 4000, Sigma 46944) at 500 mg/kg body weight. Serum was collected 4 h after gavage, and the fluorescence intensity was measured on a Molecular Devices fluorescence microplate reader.

### Isolation of Lamina Propria Cells

Single cells were isolated from small intestinal lamina propria as previously described (49). Briefly, the proximal 10 cm of small intestines from euthanized mice were flushed with cold PBS to clear feces and mucus. Excess mesenteric fat and Peyer patches were removed. Intestines were opened longitudinally and incubated in pre-digestion buffer (calcium- and magnesium-free HBSS, 3% FBS, 10 mM HEPES pH 7.4, 1 mM dithiothreitol, 5 mM EDTA) twice for 15 min each, rinse buffer (calcium- and magnesium-free HBSS, 3% FBS, 10 mM HEPES pH 7.4) once for 5 min, and digestion buffer (RPMI with 3% FBS, 10 mM HEPES pH 7.4, 0.03 mg/mL DNase I, 0.1 mg/mL Liberase TM) for 10 min, all at 37 C with agitation at 220 rpm. Enzyme-digested tissue was further dissociated in a gentleMACS C-tube using the gentleMACS Dissociator *m_intestine* program, added to RPMI with 10 mM HEPES pH 7.4 and 3% FBS, and filtered through 70- or 100-micron filters. Leukocytes were enriched from the interface of a 40/80% Percoll (GE Healthcare 17-081-01) gradient after centrifugation.

### Flow Cytometry

Single cell suspensions were twice rinsed with PBS, stained with an amine-reactive viability dye in PBS for 15 min, quenched and washed with FACS wash buffer (FWB: HBSS or PBS with 0.5% BSA), blocked with 2 micrograms FcBlock (per 1 million cells) for 15 min, and stained with antibodies against cell surface antigens for 30 min. For intracellular antigens, cells were subsequently fixed in 5% neutral-buffered formalin for 30 min at room temperature, permeabilized with eBioscience Perm Buffer (Thermo Fisher 00-8333-56), blocked with normal rat serum (STEMCELL Technologies) for 15 min, and rocked with primary antibodies overnight at 4 C. Cells were washed twice with FWB prior to analysis on a flow cytometer.

Antibodies for flow cytometry were purchased from Thermo Fisher Scientific, BioLegend, BD Biosciences, or Miltenyi Biotec. For staining mouse cell surface proteins: CD45 (30-F11), CD90.2 (30-H12), TCRb (H57-597), CD4 (GK1.5). For staining mouse intracellular proteins: IL-17A (TC11-8H4), IL-22 (poly1564), and Rorγt (B2D). For staining human intracellular proteins: RORγt (REA278). For intracellular cytokine staining, cells were first incubated for 4 h in medium with 100 ng/mL phorbol 12-myristate 13-acetate (PMA), 1 μg/mL ionomycin, and 5 μg/mL brefeldin A.

### Murine *in vitro* TH17 Differentiation

Naïve CD4 T cells were isolated from murine splenocytes using a Mouse CD4 naïve T cell negative selection kit (StemCell Technologies 19765) per manufacturer’s instructions. Isolated cells were subsequently differentiated in plates pre-coated with 2 ug/mL anti-CD3 (overnight at 4 C, or 2 h at 37 C) in TH17 medium (IMDM with 10% FBS, 50 uM beta-mercaptoethanol, 2 ug/mL anti-CD28 (BioXCell BE0015, clone 37.51), 20 ng/mL recombinant murine IL-6 (R&D Biosystems 406-ML), 5 ng/mL recombinant human TGFb1 (PeproTech 100-21), 10 ug/mL anti-IL4 (BioXCell BE0045, clone 11B11), and 10 ug/mL anti-IFNg (BioXCell BE0055, clone XMG1.2)) at a density of 1 million cells per mL. Medium was replenished on day 2, and cells were split into fresh medium on day 3 of differentiation to maintain a cell density of 1-2 million cells/mL.

### RNA Isolation and Quantitative Real-Time PCR (RT-PCR)

Total RNA was isolated from whole intestinal tissue or single cell suspensions (from epithelial fractions or lamina propria) using QIAGEN RNAeasy extraction kits or TRIzol per the respective manufacturer’s instructions. Whole tissues were lysed in either ice-cold TRIzol or Qiagen RLT Buffer using Lysing matrix D (MP Biomedical 116913050-CF) with MP Biomedical FastPrep-24 Homogenizer (1 cycle of 20 s at 4 m/s). Single cell suspensions were lysed directly in TRIzol. RNA was reverse-transcribed using High Capacity cDNA Reverse Transcription kit (Thermo Fisher 4368814) per manufacturer’s instructions and assayed by quantitative real-time PCR using Taqman probes or the SYBR Green method. Relative mRNA expression was calculated using the Livak (Delta-Delta Ct) method, using *Actb* as a loading control, unless otherwise noted. In experiments using human cells, the housekeeping genes *ACTB*, *GAPDH*, *EIF4G2*, and *RPLP0* were collectively used, and the geometric mean was used in calculating relative mRNA expression.

### Single-cell RNA Sequencing and Data Analysis

Lamina propria cells were isolated from small intestines of two mice of each genotype as described above. Isolated cells were negatively selected for T cells and ILCs (STEMCELL Technologies 19851) and stained with TotalSeq-C anti-mouse hash-tagged antibodies (BioLegend). Cells from each animal were separately hashtagged. For each genotype, a total of 60,000 cells were collected for processing using the 10X Genomics Chromium Next GEM Single Cell 5’ Kit v1.1. Sequences were aligned and processed using cellranger v7.1 using the mm10 reference genome and default parameters. Cellranger output was further processed with R version 4.3.3 and Seurat version 4.3 (50). Seurat objects were created using only genes that appeared in at least three cells. Cells were further filtered to exclude multiplets (defined as having 2 or more different hashtags) and low-quality/multiplet cells (cells with fewer than 200 detected genes, more than 2,500 detected genes, or more than 15% mitochondrial reads). Read counts were then normalized using NormalizeData. After viewing UMAP clusters, T and innate lymphocyte clusters were selected based on the presence of T/innate lymphocyte genes (*Ptprc*, *Trac*, *Trdc*, *Cd3e*, *Cd4*, *Cd8a*, *Ncr1, Klrb1b*) as well as the absence of non-T/non-innate lymphocyte genes (*Des*, *Acta2*, *Col1a2*, *Pecam1*, *Cdh5*, *Epcam*, *Cd79a, Mcpt1, Apoe*).

Cell clusters were generated using the Louvain algorithm implemented by the FindClusters function. Marker genes for each cluster were determined using the Wilcoxon test on the raw counts, implemented by the function FindAllMarkers, and clusters of cell types were additionally determined by manual inspection of the lists of cluster marker genes. Dimensionality reduction by UMAP was performed using the RunUMAP function with the 30 largest principal components. Visualization of all scRNA-seq data were generated using the Seurat package and/or ggplot2. To remove cell cycle phases as a source of clustering heterogeneity, Seurat’s CellCycleScoring function was used to score each cell and regress out the effects of cell cycle genes/phases.

### Bulk RNA-seq Library Preparation and Analyses

Total RNA was isolated from intact proximal small intestine as described above. One microgram of total RNA was depleted of ribosomal RNA (NEBNext rRNA Depletion Kit v2, NEB E7405) and subsequently fragmented, reverse-transcribed into cDNA, and amplified into barcoded libraries with NEBNext RNA Library Prep (NEB E7765) using custom barcoded Illumina-compatible primers. Libraries were pooled and sequenced on a Novaseq X Plus instrument as 150-bp paired-end reads.

Sequenced reads (40-70 million paired-reads per sample) were processed with fastp (51) for quality control and adapter sequence trimming. Trimmed reads were aligned to the *Mus musculus* mm10 genome using HISAT2 (52), and transcriptomes for each sample were assembled with StringTie2 (53) and UCSC’s mm10 genes.gtf annotation. After construction of non-redundant transcriptomes across all samples (stringtie --merge), expression counts were tabulated (stringtie -e) and used as input for downstream gene expression analyses. DESeq2 (54) was used for differential gene expression analyses.

### Chromatin Immunoprecipitation (ChIP)

In vitro-differentiated TH17 cells were processed for ChIP as previously described (55). Briefly, ten million TH17 cells were crosslinked at room temperature in 1% formaldehyde for 8 min, quenched with 125 mM glycine for 5 min, and resuspended in ChIP lysis buffer (10 mM Tris pH 8, 1 mM EDTA, 0.5 mM EGTA, 0.5% w/v N-lauroyl sarcosine, and protease and phosphatase inhibitors). Each 250-300 uL aliquot was individually sonicated in a Bioruptor Pico (5 cycles of 30 s ON/30 s OFF) to generate an average DNA fragment length of 250 bp as determined by agarose electrophoresis. Debris was spun down at 20,000g for 15 min, and the cleared chromatin supernatants were quantified and used for subsequent ChIP experiments.

Sonicated chromatin was diluted to 500 uL in ChIP lysis buffer with 1% Triton X-100, 0.1% sodium deoxycholate, 1 mM EDTA, and protease and phosphatase inhibitors. Chromatin was precleared with 10 microL protein G Dynabeads (Thermo Fisher 10003D) for 4 h at 4 C before incubating with 1 microg of primary antibodies overnight at 4 C. For anti-histone H3 (Abcam ab1791) or anti-acetyl histone H3 lysine 27 (H3K27ac, Active Motif 39133) ChIP, 3 or 10 micrograms of total chromatin was used, respectively. Protein G Dynabeads were incubated with immune complexes for 2 h at 4 C before magnetic capture. Immunoprecipitates were washed five times with 1 mL RIPA buffer (50 mM HEPES pH 7.6, 10 mM EDTA, 0.7% w/v sodium deoxycholate, 1% v/v IGEPAL-CA630, 0.5 M lithium chloride, and protease inhibitors), once with TE (10 mM Tris with 1 mM EDTA, pH 8) with 50 mM sodium chloride, and eluted in elution buffer (10 mM Tris, 1 mM EDTA, 1% w/v sodium dodecyl sulfate, pH8) at 56 C for 15 min. To reverse crosslinks, eluates were incubated overnight at 56 C. Samples were diluted two-fold in TE and sequentially digested with 80 microg of DNase-free RNase A at 37 C for 1 h, and then 80 microg proteinase K at 56 C for 1 h. ChIP DNA was purified by phenol/chloroform/isoamyl alcohol (25:24:1) and ethanol precipitation. DNA pellets were resuspended in 10 mM Tris pH 8 and assayed by quantitative real-time PCR using the SYBR Green method. Antibodies were titrated to ensure they were not limiting.

### ATAC-seq Library Preparation and Data Analysis

In vitro differentiated TH17 cells were used to generate Tn5-tagmented DNA fragments as described (56). Briefly, to isolate nuclei, cells were lysed in ATAC lysis buffer (10 mM Tris-HCl pH 7.5, 10 mM sodium chloride, 3 mM magnesium chloride, 0.1% v/v IGEPAL-CA630, 0.1% v/v Tween-20, 0.01% w/v digitonin) for 3 min on ice. The nuclei were briefly rinsed in wash buffer (10 mM Tris-HCl pH 7.5, 10 mM sodium chloride, 3 mM magnesium chloride, 0.1% v/v Tween-20) before incubating in transposition buffer (10 mM Tris-HCl pH 7.6, 5 mM magnesium chloride, 10% v/v dimethyl formamide, 0.1% v/v Tween-20, 0.01% w/v digitonin) with 100 nM loaded hyperactive Tn5 transposase (Diagenode C01070012) for 30 min at 37 C at 1000 rpm. Transposed DNA was isolated from Qiagen MinElute columns. Libraries were generated using published Illumina-compatible indexed oligonucleotides and 8 cycles of PCR amplification using NEBNext High-Fidelity 2X PCR Master Mix (NEB M0541). Libraries were purified from SPRIselect beads (Beckman Coulter B23317), and 51-bp paired-end sequences were sequenced on a NovaSeq X instrument. Sequences were aligned to the GRCm38/mm10 mouse reference genome using Bowtie2 v2.4.5, and PCR duplicates were removed by samtools v1.15 followed by Picard v2.27.1 (MarkDuplicates). Bigwig files were generated using UCSC’s bedGraphToBigWig script. DNA accessible regions and sites of differential accessibility were determined by MACS2 v2.2.7.1.

### CRISPR/Cas9 Editing of Human CD4 T cells and *in vitro* TH17 Differentiation

Naïve CD4 T cells were isolated from healthy donor PBMCs using a human CD4 naïve T cell negative selection kit (Biolegend 480041) per manufacturer’s instructions. Isolated cells were stimulated in stimulation medium (Immunocult XF serum-free medium plus 50 uM beta-mercaptoethanol, Immunocult anti-CD3/-CD28 cocktail (STEMCELL), and 50 IU/mL human recombinant IL-2 (NIH BRB Preclinical Biologics Repository)) for 2-3 days. Stimulated cells were subsequently electroporated in a total of 20 uL P3 solution (Lonza) with Cas9 ribonucleoprotein complexes (3.1 uM TruCut Cas9 v2 (Thermo Scientific) with 9 uM CRISPR guides (IDT)) using the Lonza Amaxa 4D-Nucleofector and program EH-115. Target sequences recognized by CRISPR guides are GATACGTCGGTACCGGACCG for the control/non-targeting guide and TTTGGCAATGCCAAGTGCAA and ACCCCCCCAAGCAGCGTTGC for A20^ZF7^ ablation. Electroporated cells were immediately placed into pre-warmed stimulation medium and incubated for 3 days. Genomic editing was confirmed to be at least 80% efficient by Sanger sequencing. Cells were subsequently differentiated into TH17 cells in plates pre-coated with 5 ug/mL anti-CD3 in TH17 medium (Immunocult XF serum-free medium with 50 uM beta-mercaptoethanol, 30 ng/mL recombinant human IL-6 (Proteintech HZ-1019), 2.5 ng/mL recombinant human TGFb1 (PeproTech 100-21), 10 ng/mL recombinant human IL-1β (Proteintech HZ-1164), 10 ng/mL recombinant human IL-23 (Proteintech HZ-1254), 10 ug/mL anti-IL4 (BioXCell BE0240, clone MP4-25D2), and 10 ug/mL anti-IFNγ (BioXCell BE0235, clone B133.5)) at a density of 1 million cells per mL. Cells were split and medium replenished on days 3, 6, and 8 of TH17 differentiation to maintain a cell density of 1-2 million cells/mL. Cells were harvested on day 9 of differentiation for analysis.

### Statistical Analyses

Statistical analyses were performed using Prism 10 (GraphPad Software). Pathology scores were assessed by an unpaired Mann-Whitney U test. Quantitative RT-PCRs were assessed by a two-tailed unpaired Student t-test with Welch correction, unless otherwise noted. A significance level of 0.05 was used as the threshold for statistical significance. All data shown are representative of at least two independent experiments.

## Author Contributions

C.J.B., D.S., X.S, and H.S. designed and conducted experiments. E.Y., N.L., Y.S., and R.A. provided technical assistance and conducted experiments. C.J.B., M-C.K., and C.J.Y perfomed and interpreted transcriptomic and epigenetic data. B.R. contributed mice and helpful discussions. J.T. and P.T. conducted experiments with germ-free mice. C.J.B., B.A.M., and A.M. conceptualized the overall project, acquired funding, supervised experiments, and wrote and edited the manuscript. The order of co-first authors was determined by the volume and conceptual novelty of work each contributed.

## Acknowledgments

We thank members of the Ma lab for helpful discussions and advice. We thank Claire O’Leary for advice regarding isolation of intestinal lamina propria cells. This work was supported by NIH R01 CA266755 and the Crohn’s and Colitis Foundation (A.M.). Research was supported by equipment available in UCSF’s Parnassus Center for Advanced Technologies (PCAT). Flow cytometry was performed in UCSF’s Parnassus Flow Cytometry Core (RRID:SCR_018206), which is supported in part by NIH P30 DK063720 and NIH S10 Instrumentation Grants S10 1S10OD026940-01 and S10 1S10OD021822-01. Sequencing was performed at the UCSF’s Center for Advanced Technologies (CAT), which is supported by UCSF PBBR, RRP IMIA, and NIH 1S10OD028511-01 grants.

