## Supplementary Figures for "A20’s Linear Ubiquitin Binding Motif Restrains Pathogenic Activation of TH17/22 cells and IL-22 Driven Enteritis"

**Supplementary Figure 1. Colons of wild-type and A20$^{ZF7}$ mice show no inflammation.** Bar, 250 microns.

**Supplementary Figure 2. Defining cluster identity and proliferative capacity after regression of cell cycle genes.**

(A) Dot plot of cluster-specific gene expression.

(B) Cell cycle scores of WT and A20ZF7 intestinal lymphocytes.

**Supplementary Figure 3. IL-17A and IL-22 expression and acetylation of H3K27 at extragenic ATAC-accessible loci in WT versus A20$^{ZF7}$ TH17 cells.**

(A) Gating strategy to identify live TCRβ+ CD4+ cells expressing IL-17A and/or IL-22.

(B) IL-17A and IL-22 expression by flow cytometry in murine WT or A20$^{ZF7}$ cells (differentiated in vitro in TH17 conditions in the absence/presence of 100 nM FICZ).

(C) Chromatin immunoprecipitation of H3K27ac and irrelevant IgG control at DNA accessible sites. Data shown as mean $\pm$ SEM. Statistics calculated using unpaired two-tailed Student t-test with Welch correction. All comparisons are not significant.

**Supplementary Figure 4. CRISPR/Cas9-mediated editing using a dual-guide approach results in highly efficient ablation of A20$^{ZF7}$ domain.** Two CRISPR guide RNAs targeting the human A20$^{ZF7}$ domain excise 40 nucleotides from the N-terminal half of the ZF7 domain, excising the first two (or four) zinc-coordinated cysteines and forcing a frame-shift of the downstream protein-coding sequence.

(A) Agarose-TBE gel displaying PCR amplicons of *TNFAIP3*'s ZF7 domain in T cells treated with Cas9 and CRISPR guide RNAs against control/non-targeting(NT) or dual A20$^{ZF7}$ sequences. There is a visible ~40 bp decrease in amplicon size from A20$^{ZF7}$-ablated cells.

(B) Chromatogram of Sanger sequences of amplicons generated in (A) confirm 40-bp deletion within the ZF7 domain. Underlined nucleotides indicate CRISPR target sequence (black solid line) and CRISPR PAM sequence (red dotted line). Vertical dotted line indicates predicted Cas9 dsDNA nuclease cut site. Synthego's ICE analysis of chromatograms routinely shows effective ablation in >85% of alleles.

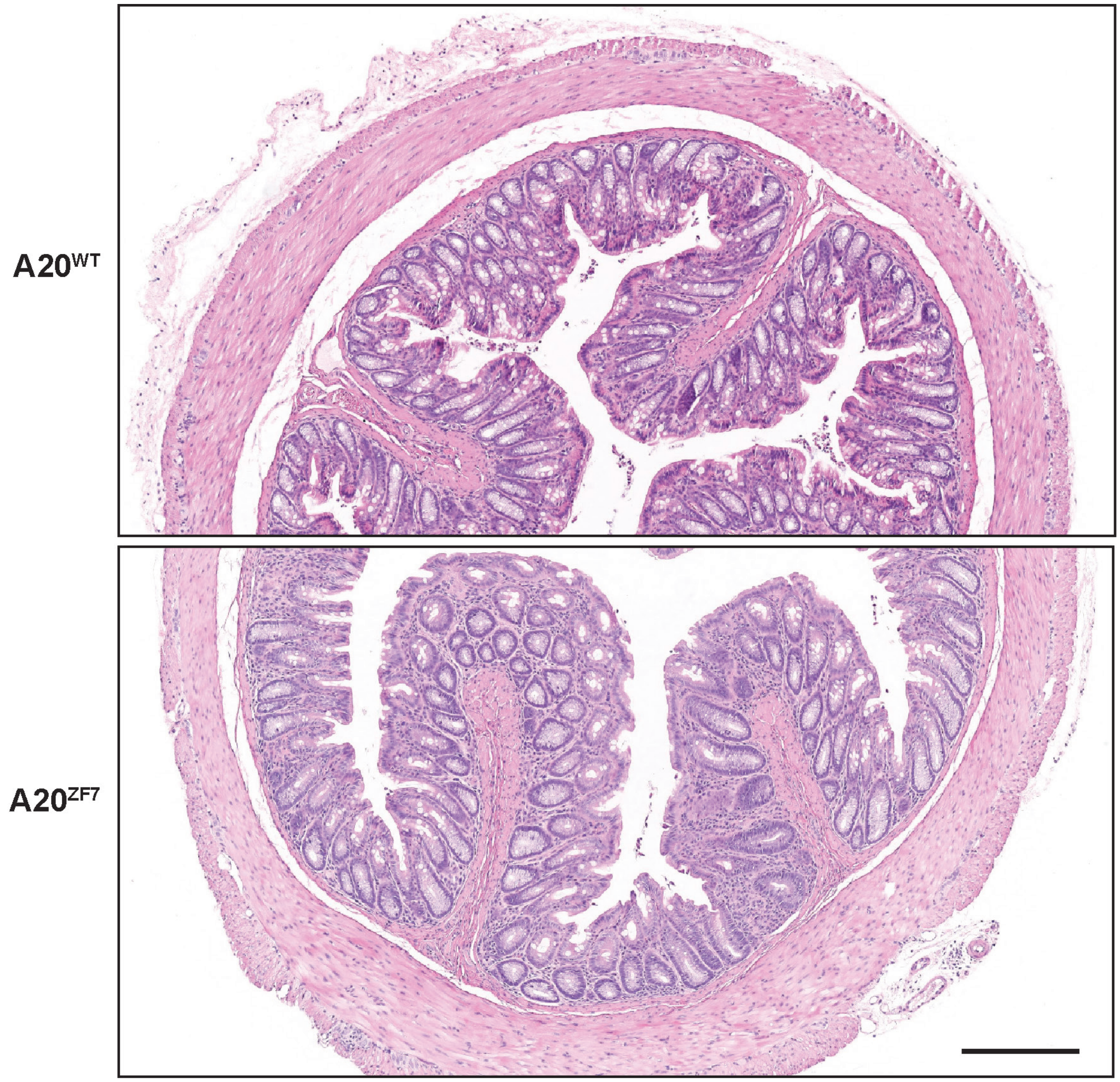


**Supplementary Figure 1.**

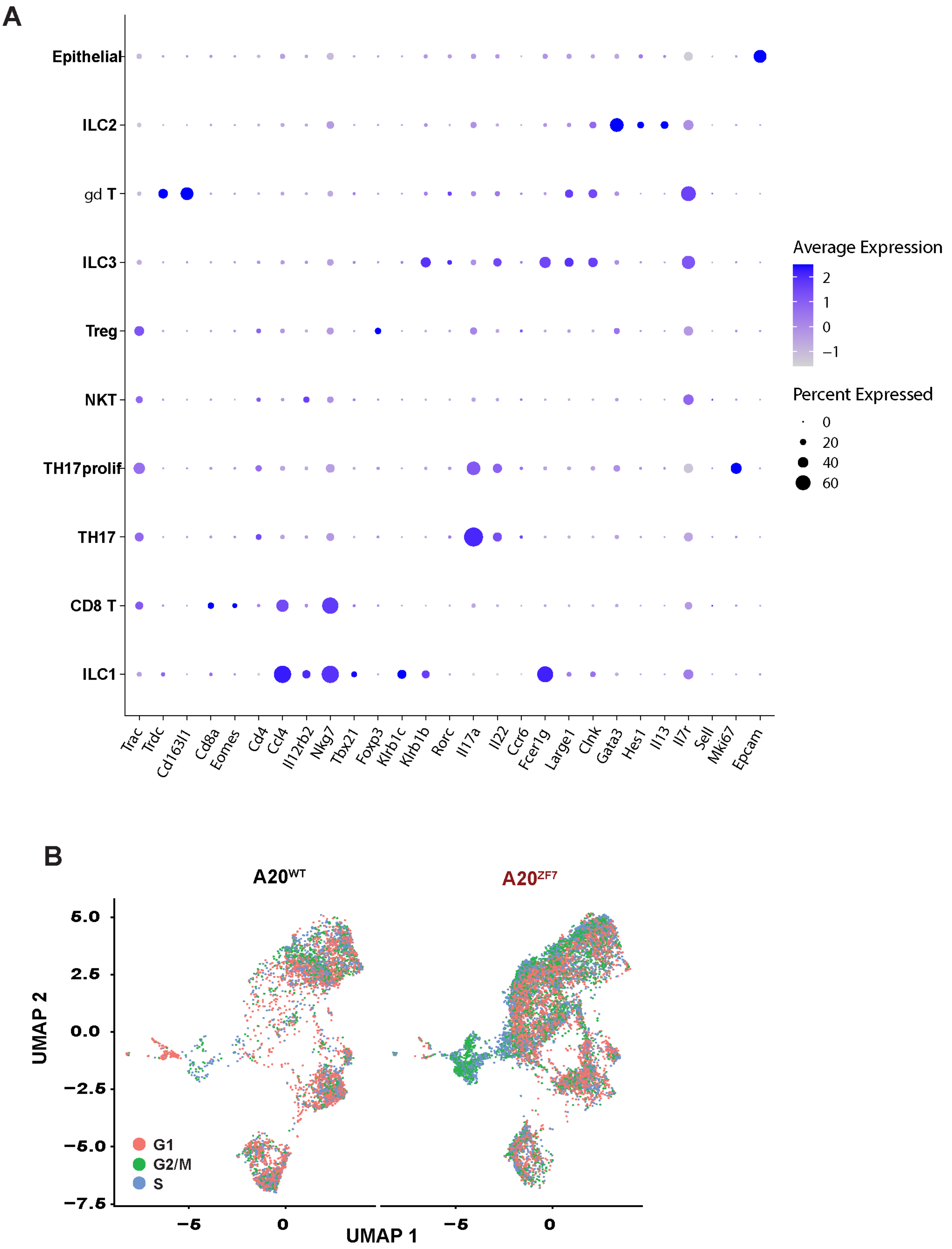


Supplementary Figure 2

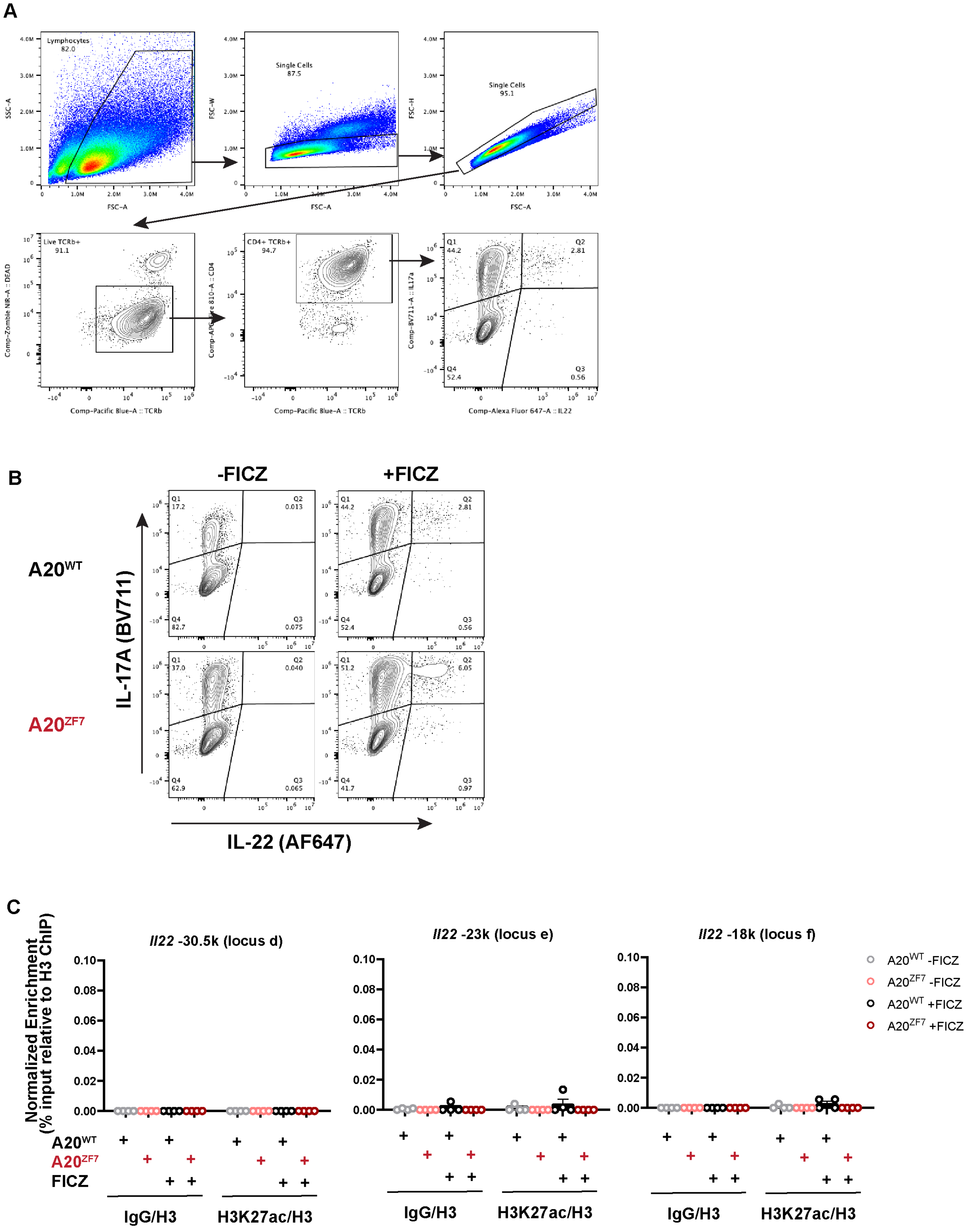


**Supplementary Figure 3**

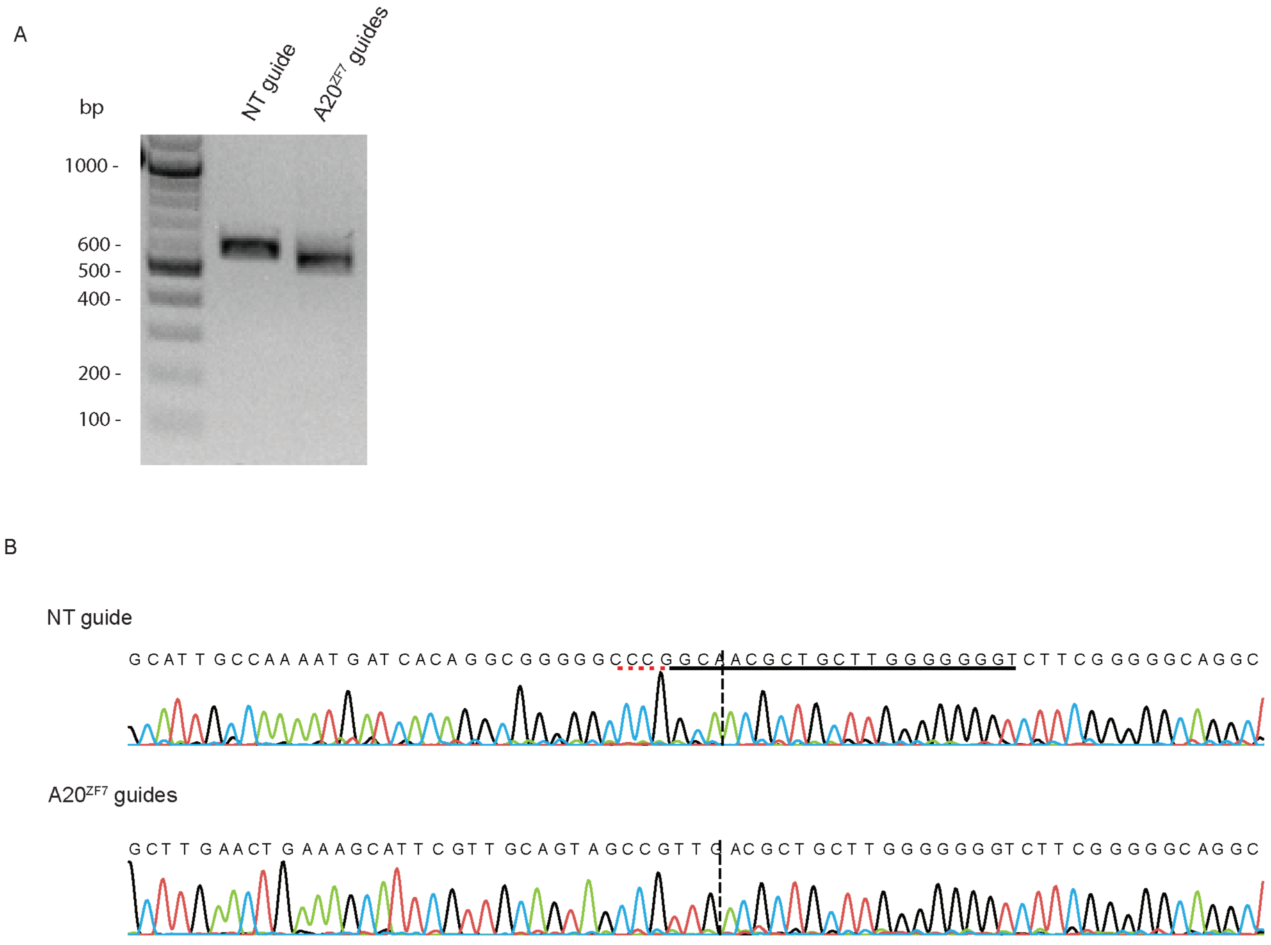


**Supplementary Figure 4**
